# Mouse Antibodies with Activity Against the SARS-CoV-2 D614G and B.1.351 Variants

**DOI:** 10.1101/2021.07.05.451203

**Authors:** Larisa Troitskaya, Nelson Lap Shun Chan, Brendon Frank, Daniel J. Capon, Brian A. Zabel, Xiaomei Ge, Dan Luo, Rachel Martinelli, Jing Jin, Graham Simmons

**Affiliations:** Hinge Bio, Inc., Burlingame, CA, 94010, USA; LakePharma, Inc., San Carlos, CA, 94070, USA; Vitalant Research Institute, San Francisco, CA, 94118, USA; Department of Laboratory Medicine, University of California, San Francisco

## Abstract

With the rapid spread of SARS-CoV-2 variants, including those that are resistant to antibodies authorized for emergency use, it is apparent that new antibodies may be needed to effectively protect patients against more severe disease. Differences between the murine and human antibody repertoires may allow for the isolation of murine monoclonal antibodies that recognize a different or broader range of SARS-CoV-2 variants than the human antibodies that have been characterized so far. We describe mouse antibodies B13 and O24 that demonstrate neutralizing potency against SARS-CoV-2 Wuhan (D614G) and B.1.351 variants. Such murine antibodies may have advantages in protecting against severe symptoms when individuals are exposed to new SARS-CoV-2 variants.

## Introduction

As the COVID-19 pandemic progresses, various resistant strains have begun to spread through the population, and it has been recognized that new antibodies will need to be added to the armamentarium in order to continue to effectively provide prophylactic and/or therapeutic value to patients. Even as more monoclonal therapeutic antibodies, identified in patients infected with severe acute respiratory syndrome coronavirus 2 (SARS-CoV-2), are authorized for emergency use, the number of SARS-CoV-2 variants that harbor mutations in the viral spike protein are increasing in incidence throughout the world^1,2^, raising questions on the long-term efficacy of current vaccines and therapeutic antibodies.

Some of the new variants of SARS-CoV-2 have been termed variants of concern (VOC) by the World Health Organization (WHO) and the Centers for Disease Control and Prevention (CDC)^3,4,5^ because they show increased transmissibility, increased virulence, or decreased effectiveness of vaccines and therapeutics^6^. These include the Alpha (B.1.1.7), Beta (B.1.351), Gamma (P.1), Delta (B.1.617.2) and Epsilon (B.1.427/B.1.429) SARS-CoV-2 variants (formerly, the United Kingdom, South Africa, Brazil, India, and California variants, respectively)^7,8^. In addition to increased transmissibility, the B.1.351, P.1 and B.1.427/B.1.429 variants have demonstrated reduced susceptibility to a combination of two therapeutic monoclonal antibodies, bamlanivimab (LY-CoV555) and etesevimab (LY-CoV016) ^2,9,10^.

Antibodies isolated from convalescent sera that neutralize the original SARS-CoV-2 isolates recognize a variety of distinct non-overlapping epitopes in the receptor-binding domain (RBD) of the spike protein^11,12,13,14.^ These antibodies, as well as antibodies from immunized individuals, frequently do not recognize at least a subset of the new strains of virus. Selection and spread of mutated disease vectors may be driven by better fitness, such as increased transmission. However, although some variants show increased transmission, as described above, many mutants do not show an improvement in fitness. It is possible that these variants are able to spread because they are not recognized by the prevalent human responses to the wild-type viruses or they have arisen by selection against the human repertoire. The difference in the murine and human antibody repertoire may allow the isolation of murine monoclonal antibodies that recognize a different or broader range of SARS-CoV-2 variants than the human antibodies that have been characterized so far.

Here, we report on the activities of two mouse monoclonal antibodies (B13 and O24), obtained from mice using the PentaMice™ platform, that recognize RBD in neutralization assays against the Wuhan (D614G) SARS-CoV-2 virus and SARS-CoV-2 pseudovirus variants. Both antibodies demonstrate excellent neutralizing potency against SARS-CoV-2 and variants we tested. B13 also binds to SARS-CoV-1. These antibodies, with their broad specificity against new variants of SARS-CoV-2 virus, provide promising candidates for therapy.

## Materials and Methods

### Cells and viruses

Vero E6 cells (ATCC) were cultured in Dulbecco’s modified Eagle’s medium (DMEM) supplemented with 10% (v/v) fetal bovine serum (FBS) and 2 mM penicillin-streptomycin (100 U/mL). 293T-hsACE2 cells (Cat# C-HA102) were purchased from Integral Molecular and cultured according to manufacturer’s recommendations. Lentiviruses pseudotyped by Wuhan D614G (Cat# RVP-702L) and B.1.351 (Cat# RVP-707L) spikes were purchased from Integral Molecular, Philadelphia, PA.

### Antibodies

LY-CoV555 (bamlanivimab), LY-CoV016 (etesevimab), AZD1061 (cilgavimab), AZD8895 (tixagevimab), VIR-7831 (sotrovimab), CT-P59 (regdanvimab), REGN10987 (imdevimab), and REGN10933 (casirivimab) were expressed in Chinese hamster ovary (CHO) cells and purified by Protein A affinity chromatography. Production of proteins was carried out by transient expression in CHO-K1 cells adapted to serum-free suspension culture (TunaCHO™, LakePharma Inc., Belmont, CA). Constructs were introduced into the LakePharma proprietary expression vector. Suspension CHO cells were seeded in a shake flask and expanded using a serum-free and chemically defined medium. On the day of transfection, the expanded cells were seeded into a new vessel with fresh medium. Transient transfections were done with the addition of the DNA and transfection reagents, under high density conditions as previously described. Transfections were carried out in cultures of 0.1 to 2.0 liters. After transfection, the cells were maintained as a batch-fed culture in a shake flask until the end of the production run. The conditioned cell culture fluid was harvested after 7-14 days, clarified by centrifugation and sterile filtered, prior to purification. The culture supernatant was applied to a column packed with CaptivA® Protein A Affinity Resin (Repligen, Massachusetts, USA) pre-equilibrated with 137 mM NaCl 2.7 mM KCl 10 mM Na2HPO4 2 mM KH2PO4 pH 7.4 (PBS). The column was washed with the PBS buffer until the OD280 value returned to baseline. The target protein was then eluted with 0.25% acetic acid buffer at pH 3.5. Fractions were collected, buffered with 1 M HEPES, and the OD280 value of each fraction was recorded. Fractions containing the target protein were pooled, formulated into 100 mM HEPES, 100 mM NaCl, 50 mM NaOAc, pH 6.0, and filtered through a 0.2 μm membrane filter and stored at 4°C prior to use. The protein concentration was calculated from the OD280 value and the calculated extinction coefficient. B13 and O24 are monoclonal antibodies isolated from the PentaMice™ platform (LakePharma Inc., Belmont, CA) after immunization with SARS-CoV-2 spike trimer protein. The B13 and O24 mouse variable (V) regions were expressed as human chimeric antibodies combining the human immunoglobulin G1 (IgG1) and kappa chain constant regions (Supplemental Figure S1).

### Live SARS-Cov-2 neutralization assay

USA-WA1/2020 (NR-52281) and Germany/BavPat1/2020 (NR-52370) were obtained from BEI Resources, Manassas, VA and expanded using a single passage in Vero/TMPRSS2. Virus was endpoint titrated in Vero/TMPRSS2 and 100 TCID50 used per well. Virus was pre-incubated for 1 hour at 37 °C with serial dilutions of antibody and then plated in replicates of 8 on Vero/TMPRSS2 cells. After 7 days wells were scored for cytopathic effect.

### Pseudovirus SARS-CoV-2 neutralization assay

The neutralization assay was carried out according to the manufacturer’s protocols. In brief, serially diluted antibodies were incubated with pseudotyped SARS-CoV-2-Renilla Luciferase for 1 hr. at 37 °C. At least nine concentrations were tested for each antibody. Pseudovirus in culture media without antibody was used as a negative control to determine 100% infectivity. The mixtures were then incubated with 293T-hsACE2 cells at 2.5×10e5 cells/ml in the 96-well plates. Infection took place over approximately 72 hrs. at 37 °C with 5% CO2. The luciferase signal was measured using the Renilla-Glo luciferase assay system (Promega, Cat# E2710) with the luminometer set at 1 ms integration time. The obtained relative luminescence signals (RLU) from the negative control wells were normalized and used to calculate the neutralization percentage for each concentration. These data were processed by Prism 9 (GraphPad) to fit a 4PL curve and calculate the log IC_50_.

### ELISA

Crossreactivity of B13 and O24 against SARS-CoV-1 spike protein was determined by ELISA. In brief, SARS-CoV-1 spike protein (UniProtKB Accession No. P59594) containing an engineered carboxyl-terminal T4 fibritin trimerization domain was expressed using the TunaCHO™ platform (LakePharma) and used to coat wells in a 384-well plate (1 μg/mL in PBS) overnight at 4°C. The wells were then washed twice (PBS with 0.05% Tween-20) and blocked (PBS with 3% BSA) for 1 h at room temperature (RT). The blocking solution was discarded, and serially diluted antibodies (3-fold dilutions from 0.001 – 200 nM) were added to the wells and incubated 1 h at RT. The plates were then washed 4 times, and then goat anti-mouse IgG-HRP (Jackson ImmunoResearch, 1:7,000 dilution in PBS with 3% BSA) was added to the wells and incubated for 1 h at RT. The plates were then washed 8 times, and chemiluminescent substrate was added (SuperSignal ELISA Pico substrate solution, Thermo, per manufacturer’s instructions). Within 15 minutes of adding substrate, the plates were read on a Molecular Devices SpectraMax M3 luminometer with Softmax Pro Version 6.2. These data were processed by Prism 9 (GraphPad) to fit a 4PL curve and calculate the log EC_50_.

## Results

### B13 demonstrates excellent neutralizing potency against SARS-CoV-2

To evaluate whether B13 can neutralize SARS-CoV-2 Wuhan (D614G) in vitro, we performed a live virus assay. Vero E6 cells were cocultured with live virus and monoclonal antibody for 7 days before measuring cytopathic effects. B13 inhibited infection of this virus with an IC_50_ value of 2.48 nM (Table 1, Figure 1).

**Table 1.** Selectivity and potency of SARS-CoV-2 monoclonal antibodies. ND = No Data

|  |  |  |  |  | Neutralization IC<sub>50</sub> (nM) |  |  |
| --- | --- | --- | --- | --- | --- | --- | --- |
|  |  | Binding |  |  | SARS-CoV-2<br>Live Virus | SARS-CoV-2<br>Pseudovirus |  |
| Antibody | Isotype | SARS-CoV-2<br>Spike | SARS-CoV-2<br>RBD | SARS-CoV-1<br>Spike | D614G | D614G | B.1.351 |
| B13 | IgG1 κ | + | + | + | 2.48 | 0.052 | 1.53 |
| O24 | IgG1 κ | + | + | - | ND | 0.024 | 0.012 |
| LY-CoV555 | IgG1 κ | + | + | - | ND | 0.028 | > 10 |
| LY-CoV016 | IgG1 κ | + | + | - | ND | 0.168 | > 10 |
| AZD1061 | IgG1 κ | + | + | - | ND | 0.035 | 0.045 |
| AZD8895 | IgG1 κ | + | + | - | ND | 0.016 | 0.088 |
| VIR-7831 | IgG1 κ | + | + | + | ND | 0.318 | 0.466 |
| CT-P59 | IgG1 λ | + | + | - | ND | 0.002 | 0.286 |
| REGN10987 | IgG1 λ | + | + | - | ND | 0.051 | 0.035 |
| REGN10933 | IgG1 κ | + | + | - | ND | 0.035 | > 10 |

**Figure 1.**
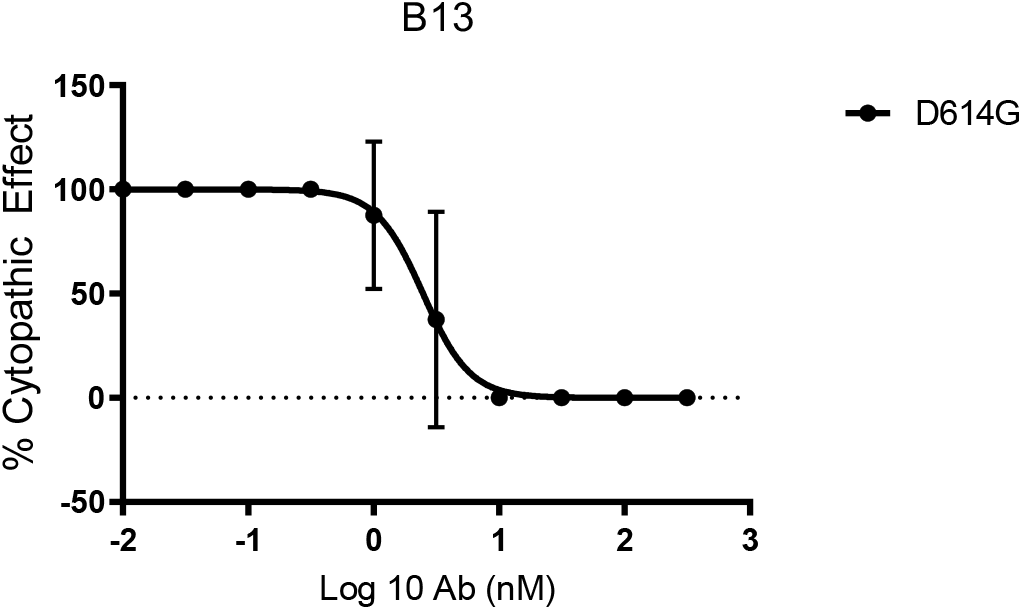
In vitro neutralization of live SARS-CoV-2. Neutralization of SARS-CoV-2 Wuhan (D614G) seven days after inoculation is shown. Values plotted are means of two replicates (n = 2), with error bars showing SD.

### O24, unlike B13, shows equivalent neutralization activity against SARS-CoV-2 and B.1.351

To assess the neutralizing efficacy of a panel of antibodies against SARS-CoV-2 Wuhan (D614G) and B.1.351 variant, we utilized a pseudovirus-based in vitro assay. 293T-hsACE2 cells were cocultured with reporter virus particles in the presence or absence of the antibodies for 72 hours before luminescence was measured (Table 1, Figure 2). B13 effectively neutralized SARS-CoV-2 D614G with an IC_50_ value of 52 pM but showed a reduced potency of 1.53 nM against the B.1.351 variant. The activity of O24 against D614G (IC_50_ = 24 pM) was comparable to that of B13. In addition, O24 had a 2-fold improvement in potency against the B.1.351 variant (IC_50_ = 12 pM) compared to D614G (Figure 2 and Table 1). While LY-CoV555, LY-CoV016, and REGN10933 were all potent against D614G (IC_50_ = 28, 170, and 35 pM, respectively), IC_50_ values against the B.1.351 variant could not be calculated because the neutralization activity was too low to detect at the concentrations tested. AZD1061, VIR-7831, and REGN10987 demonstrated comparable neutralizing potencies against D614G (IC_50_ = 35, 318 and 51 pm, respectively) and the B.1.351 variant (IC_50_ = 45, 466 and 35 pm, respectively). AZD8895 and CT-P59, while potent against D614G (IC_50_ = 16 and 2 pM, respectively), demonstrated a reduced neutralizing ability of the B.1.351 variant (IC_50_ = 88 and 286 pM. respectively).

**Figure 2.**
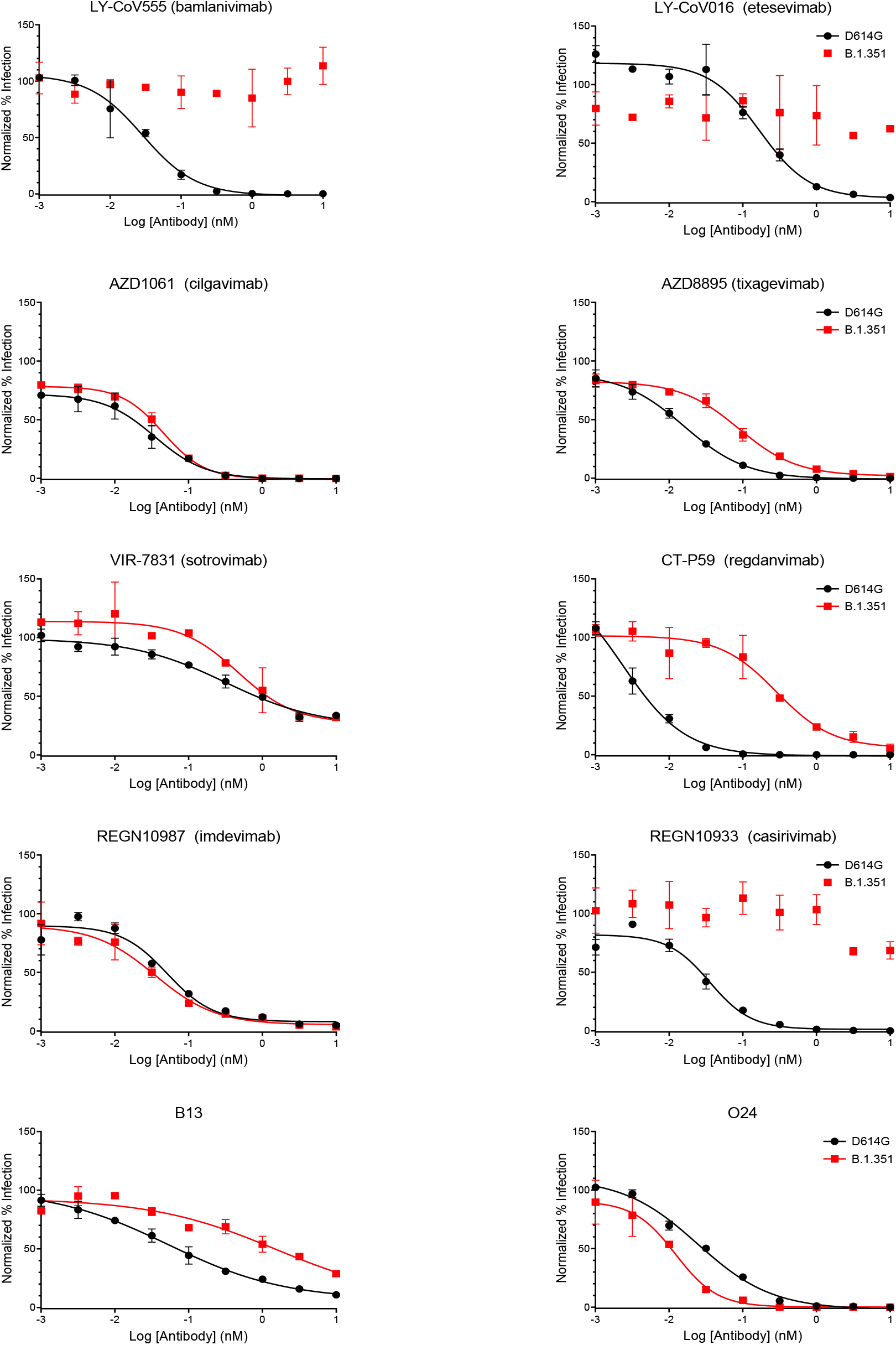
In vitro pseudovirus neutralization of SARS-CoV-2 D614G and B.1.351 variants. Neutralization of luciferase-tagged pseudotyped SARS-CoV-2 D614G (black circles) and B.1.351 variant (red squares) 72 hours after inoculation is shown. Values plotted are means of two replicates (n = 2), with error bars showing SD.

### B13 but not O24 binds to SARS-CoV-1 spike protein

B13 and O24 were evaluated for pan-coronavirus spike protein binding activity. We tested the antibodies for binding to the SARS-CoV-1 spike protein, which shares 76% identity (73% identity in the RBD domain) with SARS-CoV-2. B13 but not O24 was a potent SARS-CoV-1 spike binder, with an ELISA EC_50_ of approximately 1 nM (Figure 3).

**Figure 3.**
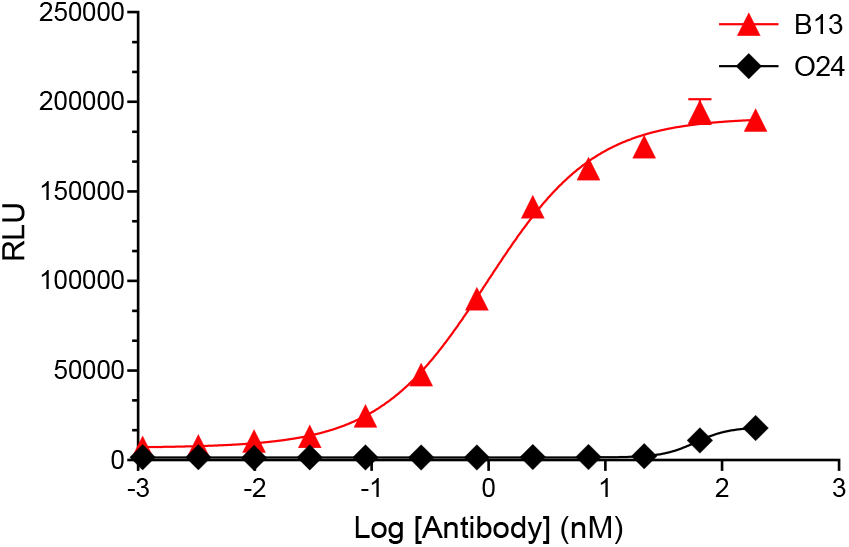
SARS-CoV-1 spike protein binding cross-reactivity. Binding of purified mouse antibodies B13 (red triangles) and O24 (black diamonds) to SARS-CoV-1 spike protein as determined by ELISA is shown. The EC_50_ for B13 binding is 0.96 nM. Values plotted are means of two replicates (n = 2), with error bars showing range. RLU: relative luminescence signal.

## Discussion

Neutralizing monoclonal antibodies targeting the SARS-CoV-2 spike protein have tremendous therapeutic potential by mitigating the symptoms of patients with mild to moderate COVID-19 or preventing infection altogether^15^. Several of these antibodies have been authorized for emergency therapeutic use as single agents or cocktails, including REGN10987 (imdevimab), REGN10933 (casirivimab), LY-CoV555 (bamlanivimab), LY-CoV016 (etesevimab), and VIR-7831 (sotrovimab). Others, including AZD1061 (cilgavimab), AZD8895 (texagevimab), and CT-P59 (regdanivimab), are currently awaiting authorization. Recently, however, due to the increase of the spread of variants, emergency use authorization of bamlanivimab as a monotherapy was revoked^16,17^ and its use in combination with etesevimab has been paused on a national basis^18^, prompting further concern about the long-term efficacy of neutralizing antibodies in development as variants continue to emerge.

Concerns have also been raised as new SARS-CoV-2 variants are emerging worldwide^2,14,19^. These new variants, especially the B.1.351 variant, clearly demonstrate antigenic drift, showing resistance to neutralizing antibodies typically derived from convalescent serum, resulting in the increase of transmissibility of SARS-CoV-2^20^. These concerns regarding resistance to neutralizing antibodies are supported by our results demonstrating the resistance of the B.1.351 variant to neutralization by LY-CoV555, LY-CoV016, and REGN10933. Interestingly, the most potent antibody against D614G, CT-P59, also displays reduced potency against the B.1.351 variant.

In order to avoid resistance to antibody neutralization by new variants, we wished to expand antibody recognition beyond the antigenic sites recognized by current human antibodies. We therefore leveraged the mouse immunoglobulin repertoire^21^ through hybridoma technology to isolate mouse monoclonal antibodies. Two antibodies, B13 and O24, were selected and identified as potent neutralizers of SARS-CoV-2 Wuhan (D614G). B13 was cross-reactive with SARS-CoV-1 spike protein (Figure 3) but demonstrated a decrease in potency against the B.1.351 variant. In contrast, O24 was not SARS-CoV-1 cross-reactive but effectively neutralized the B.1.351 variant. Our recent findings indicate that the selection of mouse antibodies such as B13 and O24 that exhibit complementary properties, may provide substantial additional coverage, particularly when engineered as intramolecular combinations, to that afforded by human-derived monoclonal antibodies against the induction of severe symptoms when individuals are exposed to new SARS-CoV-2 variants.

10-B13-A Light chain V region: DIQMTQTTSSLSASLGDRVTISCRAS<u>QDISNY</u>LNWYQQKPDGTVKLLIY<u>YTS</u>RLQSGVPSRFSGSGTGTD YSLTISNLEQEDIATYFC<u>QQGNTLPWT</u>FGGGTKLEIK

10-B13-A Heavy chain V region: QVQLQQSGAELVKPGASVKISCKAS<u>GYAFSNYW</u>MNWVKQRPGKGLEWIGQ<u>IYPGDGDT</u>NYNGNFKGK ATLTADKSSSTAYIQLSSLTSEDSAVYFC<u>GRGFIATVVEVFDH</u>WGLGTTLTVSS

10-O24-A Light chain V region: DIVLTQSPASLAVFLGQRATISCRAS<u>KSVSTSGYSY</u>MHWYQQRPGQPPKLLIY<u>LAS</u>NLESGVPARFSGSG SGTDFTLNIHPVEEEDAATYYC<u>QHSRELPPT</u>FGGGTKLEIK

10-O24-A Heavy chain V region: QAYLQQSGAELVRPGASVKMSCKAS<u>GYTFTSYS</u>MHWVKQTPRQGLEWIGA<u>IYPGNDDT</u>SYNQKFKGKA TLTVDRSSSTAYMQLSSLTSEDSAVYFC<u>TRDYSNYPFDY</u>WGQGTTLTVSS

9-K4-A Light chain V region: DIVLTQSPASLAVSLGQRATISCRAS<u>KSVSTSGYSY</u>MHWYQQKPGQPPKLLIY<u>LAS</u>NLESGVPARFSGSG SGTDFTLNIHPVEEEDAATYYC<u>QHSRELPPT</u>FGGGTKLEIK

9-K4-A Heavy chain V region: QAYLQQSGAELVRPGASVKMSCKAS<u>GYTFPNYS</u>FHWVKQTPRQGLEWIGA<u>IYPGNDDTS</u>YNQRFKGKA TLTVDKSSSTAYMQLSSLTSEDSAVYFC<u>ARDYSNYPFDC</u>WGQGTTLTVSS

5-P24 Light chain V region: DIVLTQSPASLAVSLGQRATISCRAS<u>KSVSTSGYSY</u>MHWYQQKPGQPPKLLIY<u>LAS</u>NLESGVPARFSGSG SGTDFTLNIHPVEEEDAATYYC<u>QHSRELPPT</u>FGGGTKLEIK

5-P24 Heavy chain V region: QAYLQQSGAELVRPGASVKMSCKAS<u>GYTFTSYS</u>MHWVKQTPRQGLEWIGA<u>IYPGNGDT</u>SYTQKFKGKA TLTVDKSSSTAYMQLSSLTSEDSAVYF<u>CARDYSNYPFDY</u>WGQGTTLTVSS

3-P7-A Light chain V region: DIVLTQSPASLPVSLGQRATISCRAS<u>KSVSTSGYSY</u>MHWYQQRPGQPPKLLIY<u>LAS</u>NLESGVPARFSASG SGTDFTLNIHPVEEEDAATYYC<u>QHSRELPPT</u>FGGGTKVEIK

3-P7-A Heavy chain V region: QAYLQQSGAELVRPGASVKMSCKAS<u>GYTFTSYS</u>MHWVKQTPRQGLEWIGA<u>IYPGNDDT</u>SYNQKFKGKA TLTVDKSSSTAYMQLSSLTSEDSAVYFC<u>TRDYTNYPFDY</u>WGQGTTLTVSS

10-L12-A Light chain V region: DIVLTQSPASLAVSLGQRATISCRAS<u>KSVSTSGYSY</u>LHWYQQKPGQPPKLLLIY<u>LAS</u>NLESGVPARFSGS GSGTDFTLNIHPVEEEDAATYYC<u>QHSRELPPT</u>FGGGTKLEIK

10-L12-A Heavy chain V region: QAYLQQSGAELVRPGASVKMSCKAS<u>GYTFTSYS</u>MHWVKQTPRQGLEWIGA<u>IYPGNGDT</u>SYNQKFKGKA TLTVDKSSSTAYMQLSSLTSEDSAVYFC<u>TRDYSNYPFDY</u>WGQGTTLTVSS

**Supplemental Figure S1.** Antibody V region sequences. CDR1, CDR2, and CDR3 regions for each heavy and light chain V region are underlined.

## Notes

### Competing Interest Statement

The Hinge Bio, Inc. authors own options and/or stock of the company. D.J.C. is an officer of Hinge Bio, Inc. The Lakepharma, Inc. authors are inventors on a pending provisional patent application describing some of this work.

